# Suppression of chromosome instability limits acquired drug resistance

**DOI:** 10.1101/2020.07.10.197350

**Authors:** Elizabeth A Crowley, Nicole M Hermance, Conor P Herlihy, Amity L Manning

**Affiliations:** Department of Biology and Biotechnology, Worcester Polytechnic Institute, Worcester, MA, 01609 USA

**Keywords:** Mitosis, genomic instability, Aurora B, Gefitinib, NSCLC

## Abstract

Numerical chromosome instability, or nCIN, defined as the high frequency of whole chromosome gains and losses, is prevalent in many solid tumors. nCIN has been shown to promote intra-tumor heterogeneity and corresponds with tumor aggressiveness, drug resistance and tumor relapse. While increased nCIN has been shown to promote the acquisition of genomic changes responsible for drug resistance, the potential to modulate nCIN in a therapeutic manner has not been well explored. Here we assess the role of nCIN in the acquisition of drug resistance in non small cell lung cancer. We show that generation of whole chromosome segregation errors in non small cell lung cancer cells is sensitive to manipulation of microtubule dynamics and that enhancement of chromosome cohesion strongly suppresses nCIN and reduces intra-tumor heterogeneity. We demonstrate that suppression of nCIN has no impact on non small cell lung cancer cell proliferation *in vitro* nor in tumor initiation in mouse xenograft models. However, suppression of nCIN alters the timing and molecular mechanisms that drive acquired drug resistance. These findings suggest mechanisms to suppress nCIN may serve as effective co-therapies to limit tumor evolution and sustain drug response.

**Statement of Significance:** Modulation of microtubule dynamics in cells that exhibit chromosome instability (CIN) is sufficient to promote mitotic fidelity, reduce genomic heterogeneity, and limit acquisition of drug resistance.

## Introduction

Whole chromosome instability, or numerical CIN (nCIN), is generated by underlying defects in mitosis [1, 2]. The aneuploidy that results from mitotic segregation errors promotes intra-tumor heterogeneity and is a driving force in cancer that contributes to tumor evolution and drug resistance [1–7]. Chromosome segregation is exquisitely sensitive to the regulation of dynamic microtubule attachments and defects that either increase or decrease the stability of microtubule attachments can corrupt mitotic fidelity and contribute to nCIN [8, 9]. Conversely, perturbations that reduce nCIN are proposed to limit acquired drug resistance and may hold therapeutic potential.

Aurora B kinase is a master regulator of kinetochore-microtubule dynamics during mitosis. Its overexpression is common in many cancer contexts and recent analyses of over 10,000 cancer genomes from the Cancer Genome Atlas show that, across cancer subtypes, Aurora B expression corresponds with degree of aneuploidy [10]. Increased expression of Aurora B also correlates with poor patient prognosis in a variety of cancer contexts [11]. Aurora B is a component of the Chromosome Passenger Complex (CPC). The CPC localizes to the centromere where Aurora B kinase activity regulates localization and activity of numerous kinetochore components responsible for binding and stabilizing kinetochore microtubule attachments, thereby promoting satisfaction of the spindle assembly checkpoint and regulating chromosome segregation [12]. Consistent with this function, both decreased and increased Aurora B activity at the centromere results in chromosome segregation errors [13–18].

Here we demonstrate that in a panel of non small cell lung cancer (NSCLC) cell lines, where Aurora B is highly expressed and rates of chromosome segregation are high, the experimental reduction of centromere-localized Aurora B levels, or the independent stabilization of kinetochore microtubule attachments, is sufficient to minimize genomic heterogeneity and limit whole chromosome amplifications. We additionally find that suppression of nCIN limits acquired drug resistance both *in vitro* and *in vivo,* and that sustained nCIN is rate limiting for tumor relapse.

## Materials and Methods

### Cell culture, protein expression and depletion

PC9, H1299, A549, and HBEC3-KT (ATCC) cells were grown in RPMI1640 (Gibco, #61870036) medium supplemented with 10% fetal bovine serum (Sigma, #16140071) and 1% penicillin/streptomycin (Gibco, #15070063). All cells were cultured at 37°C and 5% CO2. High resolution immunofluorescence imaging with DNA stain (DAPI) is used to monitor and confirm cell lines are free of *Mycoplasma* contamination. For short term knock down of targets, 50nM pool siRNAs (non-targeted pool or a pool of 4 siRNAs to Wapl; Dharmacon-ON-TARGETplus Human Wapl siRNA-SMARTpool) were transfected using RNAiMax (ThermoFisher, #13778100) transfection reagent according to the manufacturer’s instruction, or alternatively infected with a lentiviral construct containing a shRNA hairpin for constitutive (pLK0.1-Puro) or doxycycline inducible (tet-pLK0-Puro) depletion. Stable hairpin-expressing clones were selected with Puromycin for 7-10 days. Induced depletion was achieved by the addition of 2μg/ml doxycycline for a minimum of 48 hours.

### Immunofluorescence and Fluorescence in situ hybridization

Cultured cells or single cell suspensions derived from tumors were grown on coverslips, fixed, and stained for Aurora B (BD Biosciences, #611082), ACA (Antibodies Inc, #15-234), tubulin (Santa Cruz, #5286), CENPA S7p (Cell Signaling, #2187), and/or CENPA (Enzo Life Sciences, ADI-KAM-CC006-E) as previously described [19, 20]. To assess kinetochore microtubule stability cells were incubated in ice cold RPMI for 5 minutes (PC9 and A549) or 10 minutes

(H1299) prior to fixation and staining. Fluorescence *in situ* hybridization with centromeric probes and quantification of numerical heterogeneity was performed as previously described in [21] using alpha satellite-specific probes for chromosomes 2,6,7,8 and 10 (Cytocell, #LPE002R, 006G, 007G, 008R, 010R). For FISH from tumors, single cell suspensions were made by gently grinding tumors between two glass slides and culturing with continued drug treatment for 3-5 days. Cells were then collected and processed as above. Clonal Numerical Heterogeneity (NH) of a population was determined by scoring >300 cells per clone/tumor for copy number for chromosome probe (assessed in pairs) to determine the modal copy number, and the fraction of cells deviating from that number (i.e., the numerical heterogeneity). A given copy number with greater than 20% prevalence in a population was considered to be a stable subclone and not included in the numerical heterogeneity score. Data is represented for individual chromosomes, or alternatively, as average NH across all chromosomes measured for a given sample. A students’ 2-tailed t-test was used to determine statistical significance.

### In vivo tumor growth and relapse assays

Eight-week old male and female Crl:NU-*Eoxnf*^nu^ mice (stock #088) were purchased from Charles River Laboratory and maintained in a pathogen-free facility. All animal experiments were performed in accordance with institutional regulations after protocol review and approval by Worcester Polytechnic Institute’s Institutional Animal Care and Use Committee.

For *in vivo* growth assays, three cohorts of 5 mice each were injected subcutaneously in the flank with 5×10^6^ PC9 cells. 2 cohorts received PC9 cells expressing a tetracycline-inducible shWapl expression construct. One of these two cohorts was administered 2 μg/ml doxycycline in the drinking water. All three cohorts were monitored daily and tumor size measured 3x/week. Mice were humanly sacrificed, and tumor tissue collected when tumor volume reached 300mm^3^. For tumor relapse studies, tumor xenografts were generated as described above in 40 mice.

Xenograft tumor formation was observed in 38 mice. Once tumors reached a size of 300mm^3^ mice were randomized equally into a minus or plus doxycycline group. All mice were given 50mg/kg Gefitinib (Selleckchem, #ZD1839) by oral gavage following a 5 day on and 2 day off cycle. Mice were monitored daily and tumor size measured 3x/week to monitor tumor regression and relapse. Once relapsed tumors reached a size of 300mm^3^ mice were humanly sacrificed, tumors harvested and frozen or cultured for subsequent experimental analysis. Please see supplemental material for extended methods.

### Data Availabilitys

The data generated in this study are available within the article and its supplementary data files.

## Results

### Chromosome segregation errors in NSCLC cells correspond with high expression and enhanced mitotic centromere localization of Aurora B

Non small lung cancer (NSCLC) cells frequently exhibit high levels of whole chromosome instability (nCIN), a feature that has been experimentally linked to acquired drug resistance [1, 3–6, 22]. To understand the molecular changes that correspond with segregation errors and nCIN in NSCLC cells we first identified a panel of NSCLC cell lines that exhibit mitotic defects and chromosome copy number heterogeneity consistent with nCIN (A549, H1299, and PC9; [21]). Using immunofluorescence microscopy to assess chromosome segregation we confirm that NSCLC lines A549, H1299, and PC9 all exhibit high rates of lagging chromosomes during anaphase that are indicative of mitotic chromosome segregation errors (Figure 1A). Frequent high rates of whole chromosome segregation errors during mitosis contribute to genomic heterogeneity within a cell population that can be assessed using FISH-based approaches to measure population-level numerical heterogeneity (NH) for individual chromosomes. Using centromeric probes for chromosomes 6, 7, 8, and 10 we find that, consistent with frequent mitotic errors, populations of A549, H1299 and PC9 cells exhibit NH scores ranging from ~7-25% (Figure 1B).

**Figure 1:**
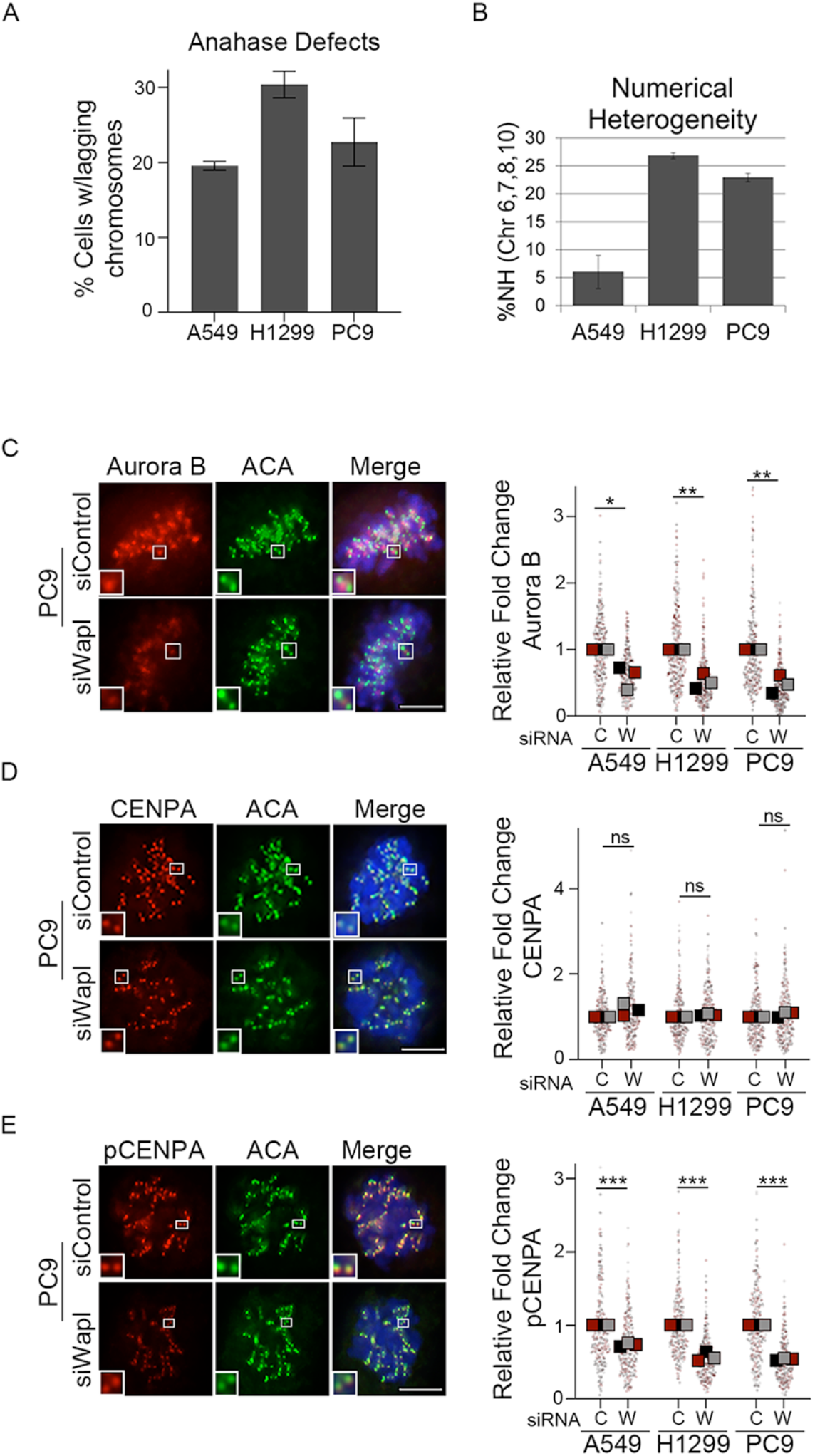
Enrichment of cohesion in NSCLC cells decreases Aurora B localization and activity at the centromere. A) Quantification of anaphase lagging chromosomes in NSCLC cell lines B) NSCLC cell populations labelled with centromere enumeration FISH probes for chromosomes 6, 7, 8, and 10 and quantified for numerical heterogeneity. A minimum of 300 cells for each chromosome probe were scored and averaged per cell line. C-E) Representation and quantification of centromere localized Aurora B, CENPA (total) and pCENPA (pSer7) in PC9 cells treated with non-targeting (siControl) or Wapl specific (siWapl) siRNA. A minimum of 90 kinetochore pairs were measured (3/cell for 30 cells), for each of 3 biological replicates. Insets are of individual kinetochore pairs at 4x magnification. Scale bars are 5μm. Here and throughout *: p< 0.05, **: p<0.01; ***: p<0.001.

Aurora B kinase, an important regulator of mitotic chromosome segregation and the mitotic spindle assembly checkpoint, is commonly overexpressed in non small cell lung cancer, where nCIN is also prevalent [23–25]. Consistent with these earlier studies, we find that Aurora B kinase is highly expressed in our panel of NSCLC cells, when compared to the normal lung epithelial cell line HBEC3-KT (Supplemental Figure 1A). Aurora B regulation of mitotic chromosome segregation is largely dependent on its localization at centromeres where it phosphorylates key substrates that regulate the stability of kinetochore-microtubule attachments [26–30] Importantly, we find that centromere localization of Aurora B in our panel of NSCLC cell lines mirrors overall expression levels of this kinase such that Aurora B localization at centromeres is increased, on average, two to five-fold over that seen in HBEC3-KT cells (Supplemental Figure 1B).

### Centromere localization and activity of Aurora B in NSCLC is sensitive to cohesion

Aurora B localization is sensitive to changes in cohesin regulation such that redistribution of the cohesin complex from pericentromere enrichment to distribution along chromosome arms leads to a concurrent redistribution of Aurora B along chromosome arms and a reduction in its centromere localization [19, 31–33] Therefore, to test the impact of high centromere Aurora B levels in the mitotic defects observed in these NSCLC lines without altering overall Aurora B protein levels, we first experimentally manipulated the distribution of the cohesin complex. Enhancement of chromosome cohesion along chromosome arms was achieved using si- and sh-RNA approaches to deplete Wapl, a well-characterized negative regulator of the cohesin complex [34] (Supplemental Figure 2A & B). Wapl associates with the cohesin complex to regulate its dynamic association with chromatin throughout the cell cycle [35] such that Wapl depletion blocks cohesin complex dissociation from chromosomes during early mitosis. We used immunofluorescence imaging of metaphase cells to identify and measure inter-kinetochore distances as a readout of functional centromere cohesion. This demonstrated that Wapl knockdown enhanced cohesion, as evidenced by a reduction in interkinetochore distance (Supplemental Figure 2C).

Using quantitative immunofluorescence, we find that depletion of Wapl perturbs Aurora B localization such that centromere-localized Aurora B levels in all three NSCLC lines is reduced by ~50% when Wapl is depleted (Figure 1C and Supplemental Figure 3A). Western blot analysis of mitotic cells indicate that this change in Aurora B localization does not arise due to Wapl-dependent changes in Aurora B expression (Supplemental Figure 2B). Consistent with reduced centromere localization and activity of Aurora B, we see comparable reduction in the phosphorylation, but not localization, of key Aurora B substrate CENPA (Total and CENPA pS7: Figure 1D & E, Supplemental Figure 3B & C).

### Modulation of Aurora B activity suppresses kinetochore-microtubule dynamics and chromosome segregation errors

Aurora B kinase is a master regulator of microtubule dynamics such that Aurora B localization and kinase activity promotes the dynamic turnover of kinetochore microtubule attachments. The fidelity of chromosome segregation has been shown to be sensitive to both increased and decreased Aurora B activity, and hence corresponding decreased and increased stability of kinetochore-microtubule attachments [8, 16, 17, 36, 37]. Consistent with this activity, we find that kinetochore fibers in Wapl-depleted cells, where Aurora B localization at centromeres had been reduced, exhibit decreased sensitivity to cold-induced microtubule depolymerization compared to control cells (Figure 2A). This suggests kinetochore microtubule attachments are stabilized by the redistribution of Aurora B away from the centromere that occurs following Wapl depletion. We next assessed mitotic fidelity in these cells and find that enhancement of chromosome arm cohesion, via Wapl depletion, is sufficient to reduce the incidence of lagging chromosomes during anaphase in both H1299 and PC9 cells (Figure 2C). Similarly, partial inhibition of Aurora B activity with the small molecule inhibitor Hesperdain, is sufficient to suppress mitotic defects in all three NSCLC lines (Figure 2C).

**Figure 2:**
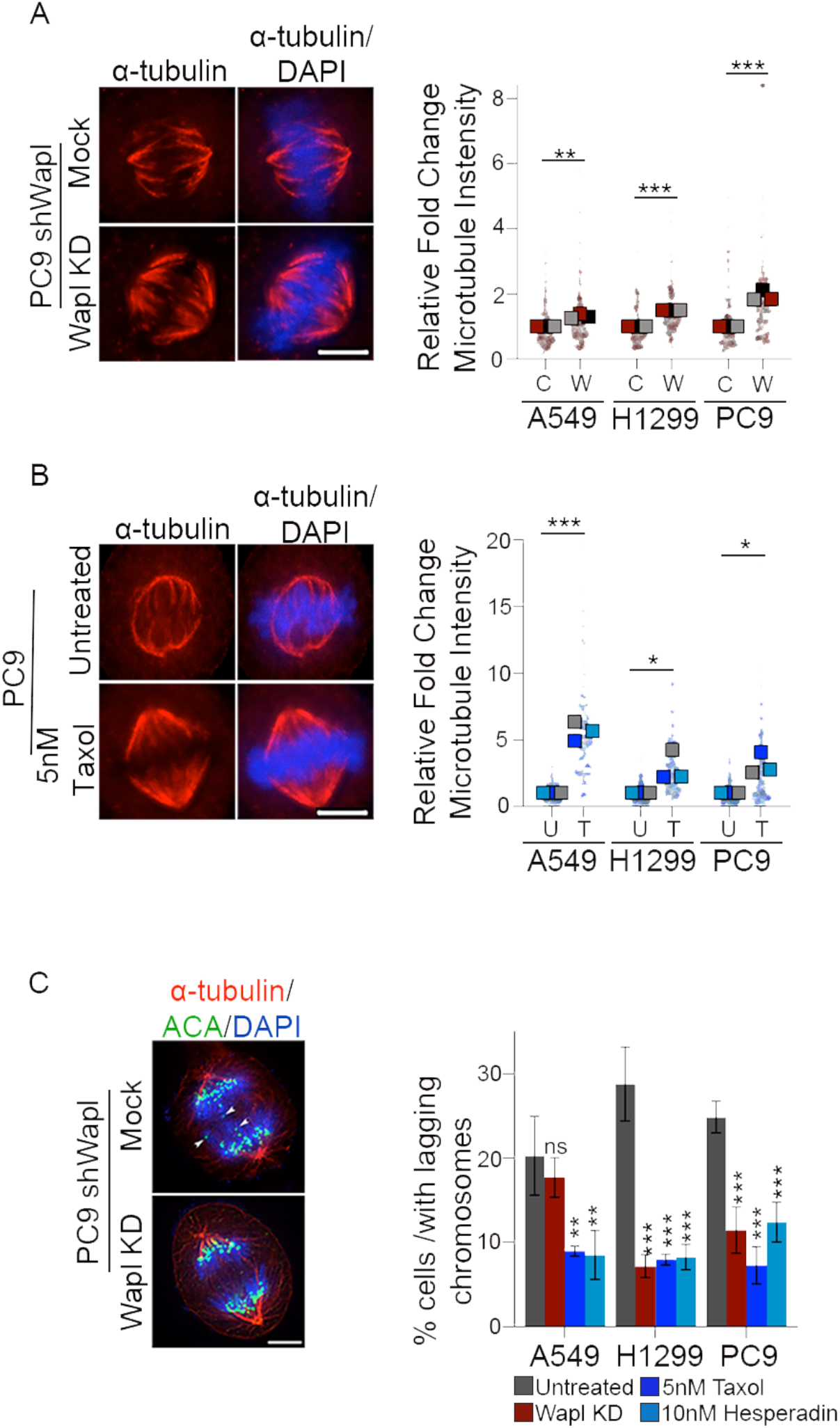
Anaphase segregation errors in NSCLC cells are reduced by microtubule stabilizing perturbations. A & B) Representation and quantification of cold stable microtubules in PC9 cells depleted of Wapl (A) or treated with 5nM Taxol (B). C) Representation and quantification of anaphase lagging chromosomes in NSCLC cells following Wapl depletion, or treatment with 5nM Taxol, or 10nM Hesperdain. White arrow heads indicate individual lagging chromosomes. A minimum of 30 metaphase or anaphase cells were scored per population for each of 3 biological replicates. Scale bars are 5μm.

To directly test the role of microtubule dynamics in the high frequency of mitotic errors observed in this panel of NSCLC cells, we treated cells with a microtubule-stabilizing concentration of Taxol prior to assessing mitotic fidelity. We find that following acute treatment (1h) with 5nM Taxol, or long-term treatment (6 weeks) with 1nM Taxol, cells exhibit both resistance to cold-induced microtubule depolymerization and reduction in anaphase lagging chromosomes, comparable to that seen following Wapl depletion (Figure 2B & C, Supplemental Figure 4A & B).

To next assess if enhanced cohesion and reduction of anaphase defects is sufficient to suppress nCIN, PC9 cells were engineered to constitutively express one of two different shRNA hairpin constructs designed to target Wapl mRNA for depletion. Single cell clones were derived from parental and Wapl-deficient PC9 cells and analyzed for numerical heterogeneity (NH). Individual clones derived from PC9 cells expressing an empty PLK0.1 vector exhibit NH values of 18-31% for chromosome 6, and up to 20-40% for chromosome 2. NH for both chromosomes was reduced ~2-3-fold in Wapl-deficient PC9 cell clones (chromosome 6: 3-12%, chromosome 2: 5-24%) (Supplemental Figure 2D), indicating that enhanced cohesion is sufficient to suppress nCIN. Together, these data support a model whereby increased Aurora B activity and/or highly dynamic kinetochore microtubule attachments underlie nCIN in NSCLC cells.

### Suppression of CIN limits drug tolerance in NSCLC cells

Activating mutations within the EGFR gene that drive tumor cell proliferation are present in nearly a quarter of all NSCLCs [38]. Patients with such mutations are commonly treated with EGFR tyrosine kinase inhibitors (TKIs) [39]. However, the majority of patients treated with EGFR TKIs ultimately develop resistance with nearly 60% of resistant or relapsed tumors exhibiting resistance-conferring mutations in the EGFR gene, making this the most frequent mechanism of EGFR TKI resistance [40]. PC9 cells exhibit an activating deletion in exon 19 of EGFR that drives cell proliferation and renders them sensitive to EGFR TKIs [41]. Like tumors in patients, these cells commonly acquire resistance to EGFR TKIs via acquisition of a secondary Threonine to Methionine mutation in EGFR (T790M) [40]. While nCIN in NSCLC, and other cancer contexts, has been correlated with acquired drug resistance [5, 23, 42], the impact of nCIN on mutation-based mechanisms of acquired drug resistance remain unclear.

Drug response is sensitive to cell proliferation rates and the impact of aneuploidies that result from nCIN have alternatively been demonstrated to promote tumor cell growth or to reduce proliferative capabilities [43, 44]. Therefore, we first assessed proliferative capacity of PC9 cells with and without constitutive nCIN/Wapl depletion (Mock and Wapl KD, respectively) or following acclimation to microtubule-stabilizing concentrations of Taxol. Importantly, in all clones tested, proliferation rates with or without Wapl depletion or Taxol treatment, when grown in the absence of TKI treatment, were comparable (Supplemental Figure 4C-E). Similarly, anchorage independent colony formation assays of growth in soft agar revealed similar colony number and size, irrespective of Wapl/nCIN status (Supplemental Figure 4F), indicating that suppression of nCIN alone does not alter drug naïve PC9 cell growth.

Next we assessed the response of cells, with and without Wapl depletion or Taxol treatment, to the EGFR TKI Gefitinib. EGFR activity results in phosphorylation of EGFR and downstream targets (such as ERK) and promotes cell proliferation. Western blot analyses indicate that PC9 cells with either mock or induced Wapl depletion or treated with 1nM Taxol are initially similarly responsive to Gefitinib: all populations show dramatic reduction of EGFR-dependent phosphorylation (phospho EGFR: Tyr1068 & phospho ERK: Tyr 202/Tyr 204) (Figure 3A). Nevertheless, following 4 weeks of continuous treatment with a sub-lethal dose of Gefitinib drug-tolerant cells slowly form colonies that can be detected with crystal violet stain. Following long-term exposure to Gefitinib, Wapl-depleted and Taxol-treated PC9 cells exhibit a dramatic reduction in the number and size of drug tolerant colonies that arise while under Gefitinib treatment (Figure 3B & C).

**Figure 3:**
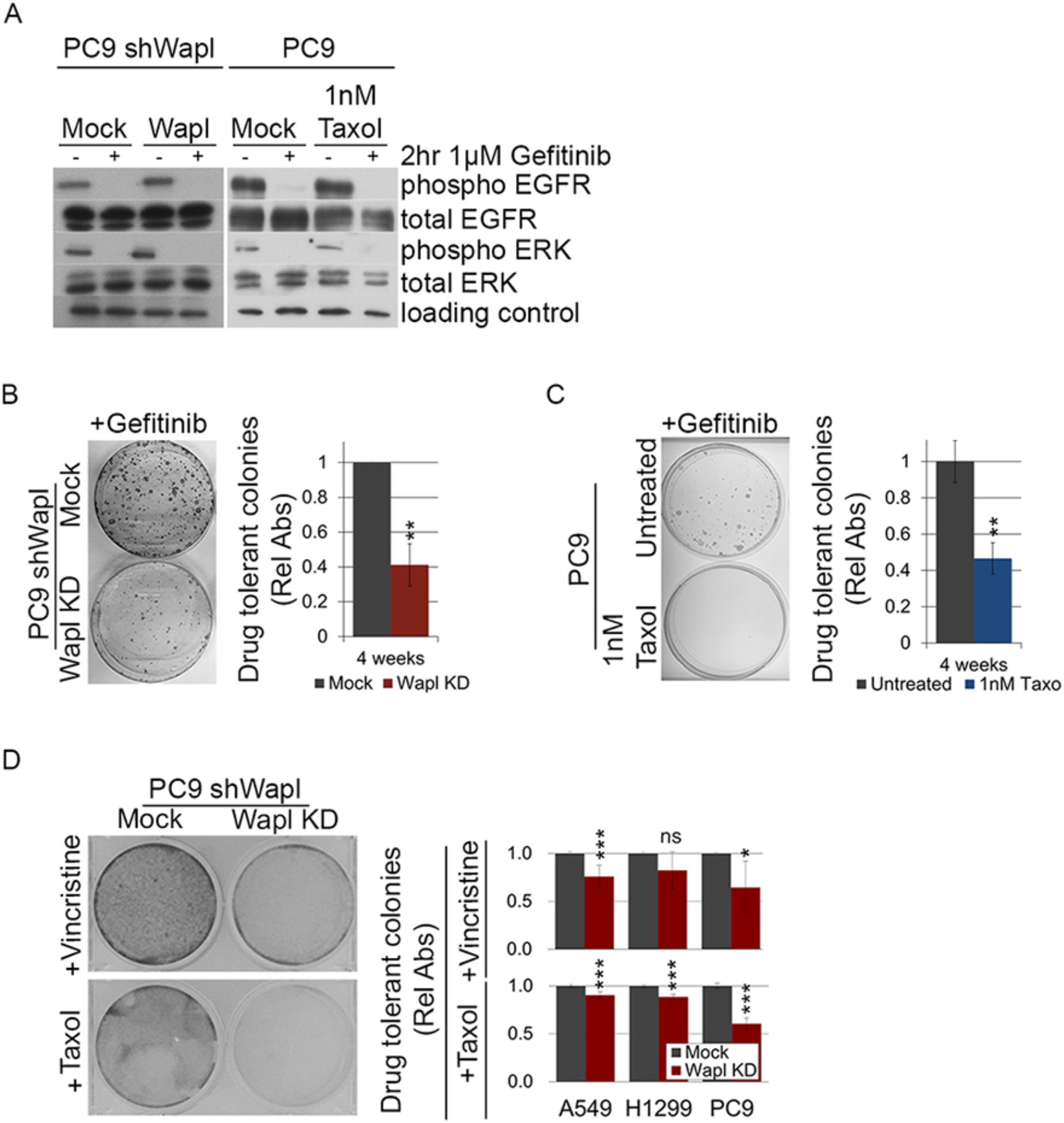
Suppression of nCIN limits drug tolerance. A) Total and phosphorylated levels of EGFR and ERK in PC9 cells with induced depletion of Wapl or 1nM Taxol treatment following Gefitinib treatment. B & C) Crystal Violet staining and quantification of drug tolerant colonies of control and Wapl depleted PC9 cells following extended culture of Wapl depleted (B) or Taxol treated (C) cells in presence of 1mM Gefitinib D) Crystal Violet staining and quantification of drug tolerant colonies of control and Wapl depleted NSCLC cells following extended culture in the presence of Vincristine or Taxol. Representative images and quantification reflect data from three independent biological replicates.

Acquired resistance to other clinically relevant therapeutics such as Vincristine and Paclitaxel have been reported in NSCLC [45–50]. Therefore, to next test whether the impact of suppression of CIN on drug tolerance is specific to Gefitinib or instead may similarly limit the outgrowth of cells tolerant to other therapeutic agents, we performed *in vitro* drug tolerance assays in A549, H1299, and PC9 cells with and without constitutive depletion of Wapl in the presence of either Vincristine or Paclitaxel. Over the course of 14 weeks the concentration of these drugs was doubled every 4 days until drug tolerant cells emerged. Comparative measures of cell numbers in each condition were approximated using crystal violet staining. Following long-term treatment with either Vincristine or Paclitaxel, all three NSCLC lines constitutively depleted of Wapl show reduction in the number of drug-tolerant cells (Figure 3D). Importantly, cells constitutively depleted of Wapl for the duration of the drug sensitivity assay show no change in proliferation rates, suggesting decreased tolerance to Vincristine or Paclitaxel cannot be accounted for by changes in proliferation rate following Wapl depletion (Supplemental Figure 4E). These data suggest that nCIN permits acquisition of drug tolerance to promote continued proliferation.

### Chromosome Instability informs mechanism of TKI drug resistance

To better understand the relationship between nCIN-dependent genomic changes that may promote or permit resistance to drug therapy we characterized four drug-tolerant clones from each of the parental and Wapl-depleted populations that were exposed to long-term Gefitinib treatment. Clones were selected and expanded under culture conditions that maintained 1μM Gefitinib and mock or induced Wapl depletion, as appropriate. Importantly, in both Wapl-depleted and Wapl-proficient contexts, drug tolerant PC9 cells that persist following long term exposure to Gefitinib continue to exhibit features consistent with Wapl status: single colonies derived from Mock-depleted cells have frequent anaphase defects and a high measure of NH and whole chromosome amplifications while Wapl-depleted cells have reduced anaphase defects and less numerical heterogeneity (Figure 4A-C).

**Figure 4:**
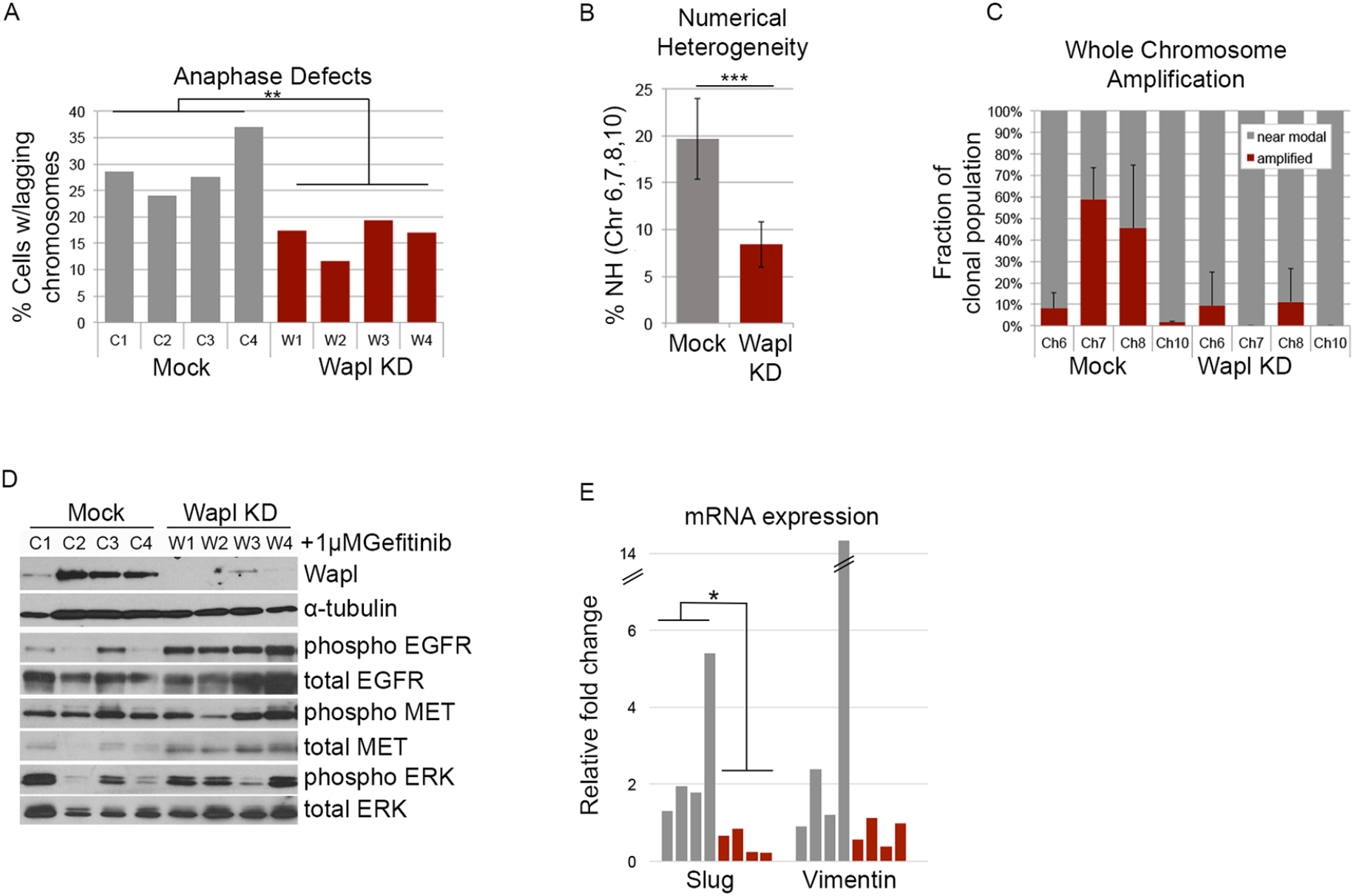
Suppression of nCIN influences mechanisms of drug tolerance in PC9 cells. A) Quantification of anaphase cells exhibiting lagging chromosomes in clonal populations of Gefitinib-tolerant cells with (Wapl KD) or without (Mock) induced Wapl depletion. B & C) Gefitinib-tolerant PC9 clonal populations derived with or without Wapl depletion labelled with centromere enumeration FISH probes for chromosomes 6, 7, 8, and 10 and quantified for chromosome copy number. Represented as average numerical copy number across all chromosomes assessed (B) and fraction of cells with modal or amplified copies of the indicated chromosomes (C). A minimum of 300 cells per condition per centromere probe assessed were scored. D) Western blot showing Wapl levels and EGFR pathway status in four each of mock and Wapl-depleted Gefitinib-tolerant clones. E) qPCR analysis of mRNA levels indicative of EMT pathway activation (Slug, Vimentin) in Mock and Wapl-depleted Gefitinib-tolerant clones.

Cells that acquire tolerance or resistance to TKI activity do so by restoring or bypassing EGFR kinase function to activate downstream pathway components and promote proliferation [40]. Although the emergence of drug tolerant clones is decreased significantly by Wapl depletion, molecular characterization of EGFR pathway function indicates that drug tolerant clones that do arise from both Mock- and Wapl-depleted populations similarly exhibit phosphorylation of ERK, a downstream target in the EGFR pathway (Figure 4D). In contrast, Wapl-depleted PC9 clones, but not Mock-depleted PC9 clones, exhibit EGFR auto phosphorylation (Figure 4D). This phosphorylation is dependent on EGFR kinase activity and suggests drug-tolerant clones that lack nCIN (Wapl KD) primarily arise through the acquisition of resistance-conferring mutations that impair TKI binding, while clones that sustain nCIN (Mock) likely exploit other mechanisms of drug tolerance [38–40, 47].

Alternative pathways reported to promote TKI resistance in NSCLC include amplification of EGFR, amplification of the receptor tyrosine kinase MET [47], and/or activation of epithelial to mesenchymal transition (EMT) [51–53]. To understand if these alternative pathways could explain tolerance of Mock-depleted PC9 clones to Gefitinib treatment, we first assessed copy number of chromosome 7, on which both EGFR and MET genes are located. Drug naïve PC9 cells have a modal copy number of four for chromosome 7, as do drug tolerant Wapl-depleted clones. In contrast, nearly 60% of drug tolerant Mock-depleted cells have >8 copies of chromosome 7 (Figure 4C, Supplemental Figure 5). We next looked at expression of key EMT transcription factor SLUG and found it to be upregulated 2-fold in 3 out of 4 Mock-depleted clones, but not in any of the Wapl-depleted clones (Figure 4E). The gene that encodes for SLUG, SNAI2, is located on chromosome 8p, adjacent to the centromere. In support of a model whereby CIN influences the mechanism of TKI resistance, single cell analysis of chromosome 8 copy number indicates that the same 3 Mock-depleted clones with increased SLUG expression also exhibit amplification of chromosome 8 (Figure 4C, Supplemental Figure 5). Consistent with SNAI2 amplification and increased Slug expression, Vimentin, a widely used marker of EMT and a Slug target gene is also increased 2 to 12-fold in Mock-depleted clones (Figure 4E). Together these data suggest that nCIN influences the mechanism of acquired drug resistance. Without nCIN, mutation-driven mechanisms of drug resistance dominate, but the acquisition of such mutations are slow, particularly in drug treated populations with limited proliferation, and as a result drug resistant colonies are slow to emerge. Through whole chromosome gains and losses hundreds to thousands of genes may become mis-regulated in a single cell division. In this way nCIN has the capacity to promote cellular changes that contribute to drug tolerance.

### Wapl depletion is sufficient to suppress nCIN in vivo

To assess how cohesion-dependent suppression of nCIN impacts tumor initiation and growth *in vivo,* PC9 cells with or without a tetracycline-inducible shWapl construct were injected subcutaneously into the flanks of nude mice to generate 3 cohorts: one with parental PC9 cells, and two with PC9 cells that harbor the inducible shWapl construct (Supplemental Figure 6A). One cohort of PC9 shWapl mice received doxycycline in the drinking water to induce expression of the Wapl-targeting hairpin throughout the duration of tumor initiation and growth. The cohort injected with the PC9 parental cell line similarly received doxycycline in the water as a negative control. Tumor initiation was comparable in all three cohorts of mice, as was rate of tumor growth (Supplemental Figure 6B, Supplemental Figure 7A). To confirm the efficiency of Wapl depletion and the suppression of nCIN in response to doxycycline treatment *in vivo,* tumors from mice in each cohort were excised when they reached 300cm^3^ and subjected to analysis. Quantitative PCR analyses confirm that depletion of Wapl mRNA and protein levels were achieved and sustained during tumor initiation in response to doxycycline (Supplemental Figure 6C). Consistent with *in vitro* assays, cells derived from Wapl-deficient tumors exhibit fewer anaphase defects (Supplemental Figure 6D) and less intra-tumor chromosome numerical heterogeneity (Supplemental Figure 6E). Together these data indicate that suppression of nCIN in NSCLC cells is not in itself sufficient to limit tumor initiation and growth *in vivo.*

### nCIN is a driver of drug resistance *in vivo*

To test the role of nCIN in tumor relapse, the xenograft model described above was used to establish mice that harbored inducible PC9 shWapl tumors. Once tumors reached a volume of 300mm^3^, mice were put on a 5 days on, 2 days off (5+2) regimen of 50 mg/kg Gefitinib treatment and randomly assigned to two cohorts. One cohort received doxycycline in the drinking water to induce Wapl depletion concurrent with Gefitinib treatment, the other did not (Figure 5A). Both cohorts of mice exhibited similar initial response to Gefitinib, with all tumors exhibiting >50% recession within 2 weeks. In the 100 days following initial tumor recession mice were sustained on a 5+2 drug regimen. During this time 37% (7/19) of Mock-depleted tumors and 21% (4/19) of Wapl depleted tumors relapsed. Relapse of Wapl depleted tumors was delayed by nearly 3 weeks compared to those without induced Wapl depletion (66 days vs 46 days for Mock depleted tumors) (Figure 5B, Supplemental Figure 7B). A similar frequency of the TKI resistance-conferring EGFR T790M mutation was detected in relapsed tumors from both cohorts (Figure 5C). Interestingly, anaphase defects were similarly prevalent in all relapsed tumors, regardless of cohort, indicating a selective pressure to maintain CIN during acquisition of drug resistance (Supplemental Figure 7C). Consistent with this, all residual/non-relapsed tumors analyzed from the Wapl depleted cohort sustained Wapl depletion while tumors that relapsed expressed Wapl at levels comparable to the Mock-depleted cohort (Figure 5D). These data support a model whereby re-establishment of Wapl expression/nCIN, or expansion of clones that fail to silence Wapl/sustain nCIN, is limiting for acquired drug resistance and tumor relapse (Figure 6).

**Figure 5:**
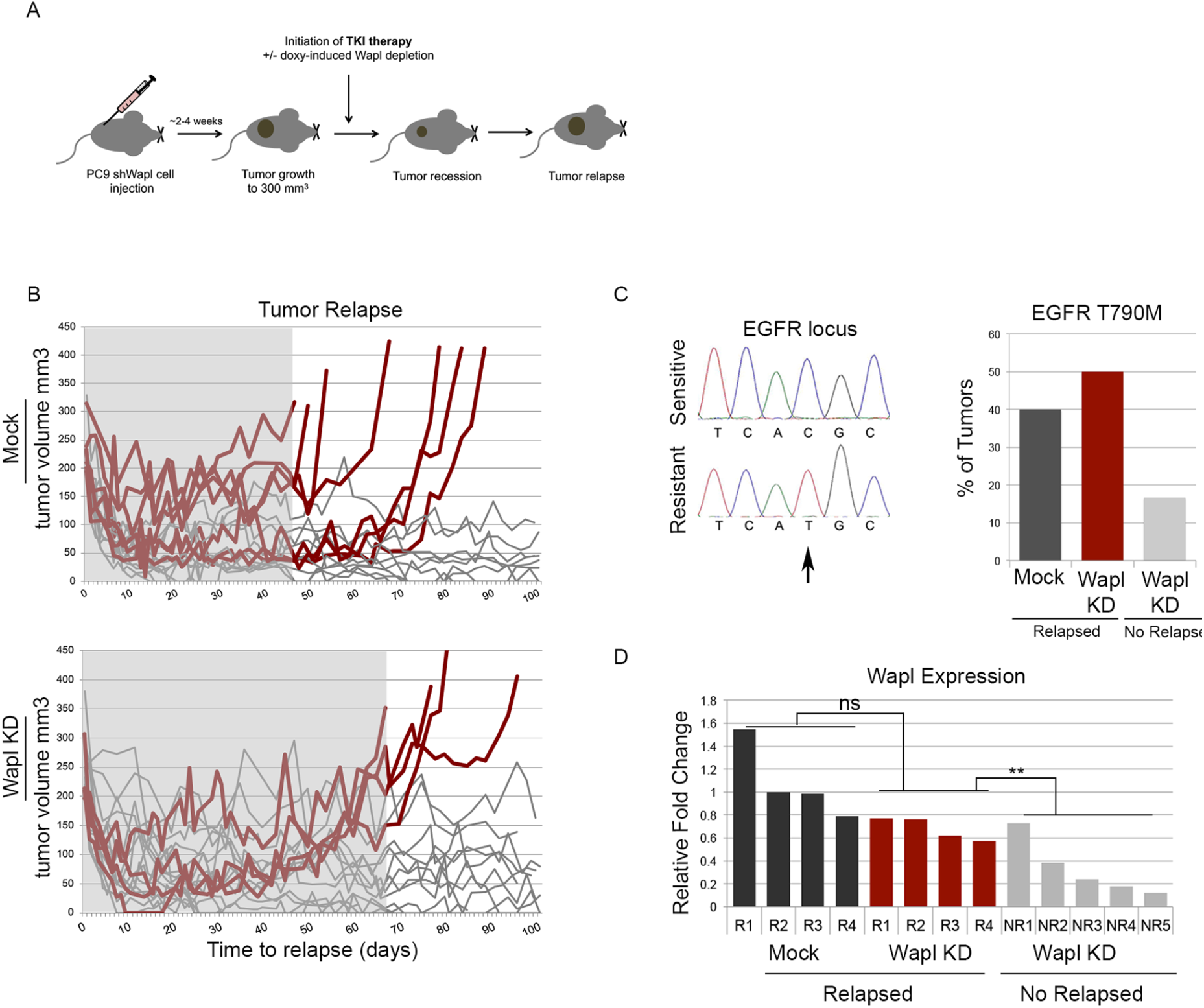
nCIN is a driving force for acquired drug resistance. A) Diagram of experimental set up: PC9 cells carrying an inducible shWapl construct were injected into the flank of the mice. Once tumors reached 300mm^3^ mice were split into two cohorts: one was administered doxycycline in drinking water to induce Wapl depletion concurrent with Gefitinib via oral gavage, the other received Gefitinib alone. All mice were monitored for initial tumor recession and relapse. B) Time to relapse of tumors with uninduced (n=19) and induced Wapl depletion (n= 19). Red lines indicate tumors that relapsed in this time frame, gray lines indicate those that remained responsive to treatment. Shaded box indicates the minimum relapse free response to Gefitinib in each cohort. C) Visualization and quantification of C à T mutation that results in the T790M mutation in EGFR that confers resistance to TKI in tumors from each cohort. D) qPCR analysis of Wapl levels in control (Mock) and induced Wapl depletion (Wapl KD) that relapsed or not following Gefitinib treatment.

**Figure 6:**
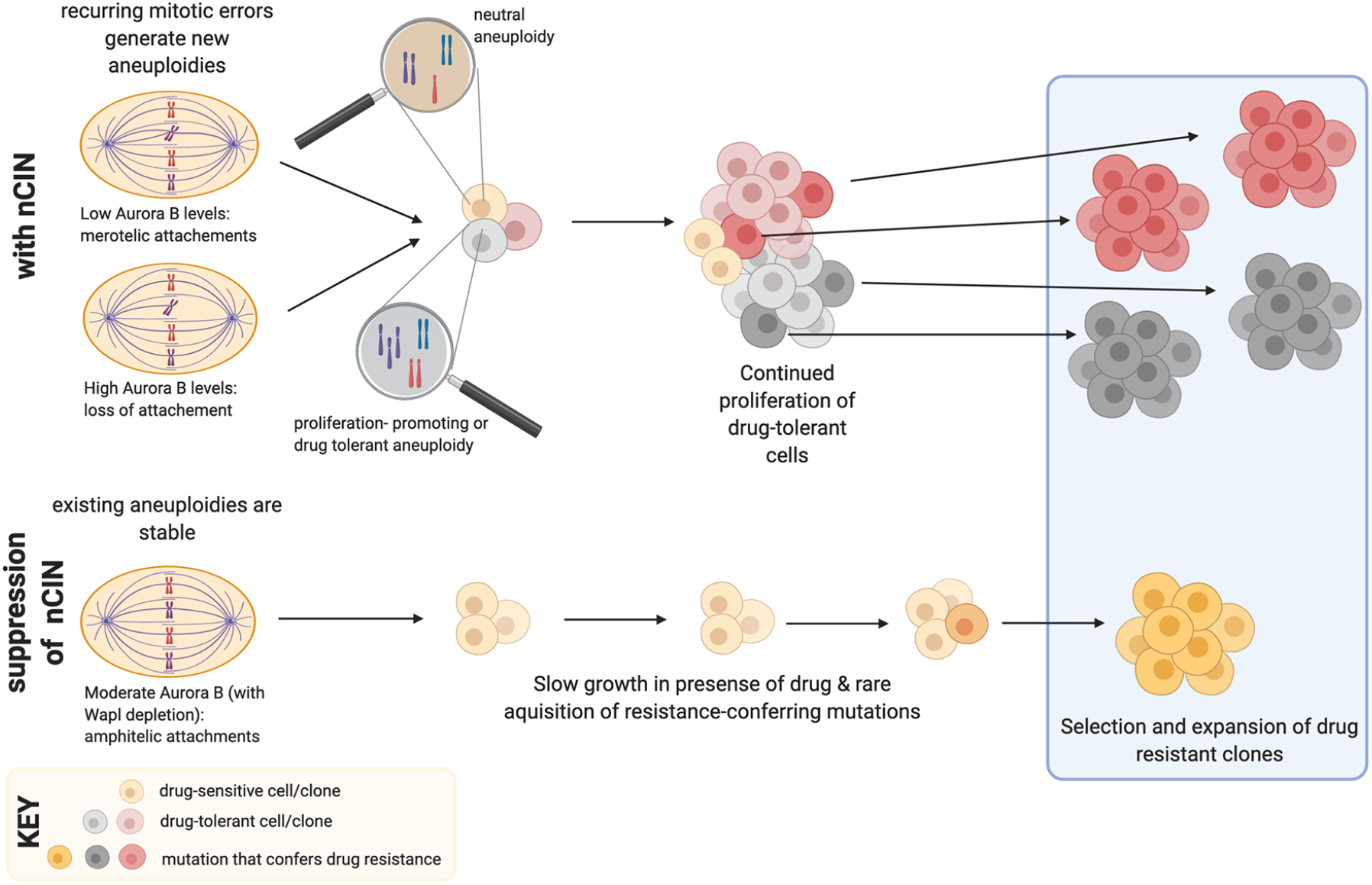
Mitotic defects that promote CIN enable generation of drug tolerant aneuploidies that permit continued growth and increased incidence of acquired drug resistance.

## Discussion

Clonal mutations that enable resistance to targeted or chemotherapeutic approaches pose a clinical challenge and remain a major cause of death in many cancer types [54]. Clinically and experimentally, the degree of intra-tumor genomic heterogeneity, and underlying defects in mitotic cell division have been functionally linked to tumor evolution, drug resistance, and metastasis [3, 4, 6, 7, 55, 56]. In the context of NSCLC, identification of driver mutations in EGFR and the initial clinical success of TKI treatment has been hampered by rapid and prevalent acquisition of drug resistance [38]. Here, we present mechanistic evidence that nCIN may arise from mis-regulation of cohesin-sensitive Aurora B kinase activity at centromeres and deregulation of spindle microtubule dynamics. We propose that subsequent chromosome amplifications contribute to a high incidence of acquired resistance to therapy. The reduction in emergence of drug tolerant clones following experimental suppression of nCIN demonstrate that whole chromosome copy number changes create a favorable environment for continued proliferation while resistance-conferring mutations are attained. Consistent with this, we find that a selective pressure exists *in vivo* to maintain or re-establish nCIN to facilitate robust acquisition of drug resistance and tumor relapse (Figures 5 & 6).

### Whole chromosome segregation errors enable tumor evolution and drug resistance

Our analysis examined isogenic PC9 cells that primarily differ, at least initially, in their nCIN status. Our results indicate that nCIN contributes to drug resistance by allowing for the generation of whole chromosome amplifications that harbor key drug tolerance genes (like EGFR, MET, and SNAI2/SLUG) and promote continued proliferation in the presence of TKI. In the absence of an increased mutation rate, this continued proliferation is key to enable the acquisition of replication-dependent mutations that confer robust drug resistance and tumor relapse. Such adaptive mutability may be particularly relevant to the mis-segregation and subsequent selection for amplification of chromosome 7, which contains both MET and EGFR genes. Increased EGFR gene copy number both promotes proliferation [40] and, by virtue of having more EGFR gene templates for replication-acquired mutation, increases the apparent mutability of individual cells [57]. These findings are consistent with previous studies showing selective pressure for cancer cells to sustain nCIN and demonstrating that high levels of nCIN can drive the generation and selection of drug-resistant clones [6, 7, 58, 59].

### High mutation rates may preclude the need for nCIN in acquired drug resistance

In the absence of nCIN, resistance is limited to the clonal amplification of cells with pre-existing amplifications or mutations, and those that acquire chromosome amplifications through rare segregation errors. An increase in mutation rate may negate the need for nCIN by increasing the frequency at which resistance-conferring mutations are generated in each cell cycle. Consistent with this view, high mutation rates and nCIN have been found to be mutually exclusive in various cancer contexts [60].

### nCIN as a therapeutic target

Similar to what we demonstrate here, work from other groups has shown that presence of nCIN can promote acquisition of drug resistance and is a mechanism to evade oncogene addiction [5, 58]. Our data additionally show that suppression of segregation errors in a cancer context is achievable and that reduction of nCIN can limit mechanisms of acquired drug resistance. Indeed, due to its role in regulation of chromosome segregation, Aurora B is a provocative drug target, and its inhibition has recently been shown to be efficacious in limiting proliferation of TKI-resistant NSCLC cells [61]. Together these data propose that pathways that promote nCIN may serve as valuable drug targets, alone, or as co-therapies to enhance or prolong response to targeted therapeutic approaches.

## Supporting information

supplemental methods and figures with legends

## Acknowledgements

This work was supported by a Smith Family Award for Excellence in Biomedical Research and R00CA182731 to ALM.

## Author Contributions

ALM conceived and devised the study. ALM, EAC, and NMH designed the experiments. ALM,

NMH, EAC, and CPH performed the experiments. ALM wrote the manuscript with input and approval from all authors.

