## supplemental methods and figures with legends for "Suppression of chromosome instability limits acquired drug resistance"

**Confirmation of protein expression and depletion**

For quantification of mRNA expression levels for Wapl, Aurora B, Slug, Vimentin, and/or expression in cell lines, drug tolerant clones, or tumors following Gefitinib treatment with or without Wapl depletion, RNA was extracted from cell lines or frozen tumors using Trizol reagent (Invitrogen, #15596026) according to the manufacturer's instructions. Complementary DNA was synthesized from 2μg of total RNA using Superscript First-Strand Synthesis System (Invitrogen, #18091050) and random hex primers. Gene expression was determined by using the ΔΔ cycle threshold method normalized to either *GAPDH* or *β-actin.* Primer sequences used for qRT-PCR can be found in Supplemental Table 1. Samples were run as technical duplicates and each experiment performed in biological triplicate. Data presented represents average and standard deviation between biological replicates. A students' 2-tailed t-test was used to determine statistical significance.

To assess protein levels, whole cell extracts were prepared using 2x Laemmli buffer (BioRad, #1610737) with β–Mercaptoethanol (Sigma Aldrich, #444203). Protein concentrations were normalized to total cell number and samples run on an SDS-PAGE gel. Proteins were transferred to PVDF membrane (Millipore) and membranes blocked in 1xTBST supplemented with 5% milk powder. Antibodies were diluted in 1xTBST/5% milk (phospho EGFR (Cell Signaling, #3777S), total EGFR (Cell Signaling, #4267S), total MET (Cell Signaling, #3127), total ERK (Cell Signaling, #9102L), WAPL (Bethy, #A300-268AI), Aurora B (BD Biosciences), Histone H3 (Abcam, #176842), and alpha-tubulin (Santa Cruz)) or 1xTBST/5%BSA (phospho MET (Cell Signaling), phospho ERK (Cell Signaling, #9101S) and incubated at 4° overnight. Membranes were washed in 1xTBST and incubated 1h in corresponding secondary antibody (rabbit, mouse: GE Healthcare, #NA391), washed in 1xTBST buffer and developed using ProSignal Pico (Prometheus, #20-300B).

**Immunofluorescence**

Images were captured using a Zyla sCMOS camera mounted on a Nikon Ti-E microscope, with a 60X Plan Apo oil immersion objective and capturing 0.3 μm z-stacks. To assess centromeric Aurora B, CENPA S7p, and CENPA levels, NIS-elements Advanced Research software was used to perform line scans in a single focal plane through individual ACA-stained kinetochore pairs where interkinetochore distance is represented by the distance between ACA peaks, and the area under the curve in the Aurora B, CENPA S7p, or CENPA stained channel indicates centromere/kinetochore-localized staining. Kinetochore microtubule intensity was assessed by measuring the intensity across a line positioned parallel to metaphase-aligned chromatin. For interkinetochore distances and Aurora B, CENPA S7p, and CENPA localization a minimum of 3 kinetochore pairs per 30 cells, per condition (90 kinetochore pairs/condition) were measured in each of 3 biological replicates. Anaphase defects and kinetochore microtubule intensity were assessed in a minimum of 30 cells per condition, in each of 3 biological replicates. Data presented represents average and standard deviation between biological replicates. A students' 2-tailed t-test was used to determine statistical significance. For figure generation, images were prepared using NIS Elements deconvolution software and represented as projections of 5 central plains. Insets represent a single focal plane.

**Growth and survival analyses**

Proliferation rates of parental cell lines and/or cells stably expressing a doxycycline inducible construct to target Wapl for depletion were plated following presence or absence of 2 μg/ml doxycycline for 2 days to assess acute depletion or for 14 weeks to assess sustained depletion. Following depletion of Wapl, cells were collected and counted for 5 consecutive days and growth rates calculated. Growth rates were similarly assessed following exposure of PC9 cells to 1nM Taxol for 6 weeks. Anchorage-free proliferation assays were performed in PC9 cells with or without constitutive shRNA-depletion of Wapl. $4 \times 10^4$ cells were mixed with 0.4% agarose in growth medium (RPMI 1640 supplemented with 10% FBS, 1% penicillin/streptomycin), plated in a 6 well dish containing a solidified layer of 0.5% agarose in growth medium and placed at 4° for 15 minutes to allow solidification. Cells were fed once a week with 2% agarose in growth medium. At 4 weeks colonies were imaged and counted. Data presented represents average and standard deviation between three biological replicates. A students' 2-tailed t-test was used to determine statistical significance.

For *in vitro* drug tolerance assays $1 \times 10^4$ PC9 shWapl cells were plated in a 10 cm dish with or without 2μg/ml doxycycline. Cells were then treated with 1μM Gefitinib (Selleckchem). Growth medium containing drug was replaced twice per week. At 4 weeks cells were washed in 1xPBS and fixed and stained with 0.5% (w/v) crystal violet (acros organics, #548-62-9) in 25% methanol (v/v) (Sigma, #67-56-1) for 30 min, rinsed 3 times with water, and dried. To quantify the relative number of drug tolerant cells, crystal violet was resuspended in 10% (v/v) glacial acetic acid and absorbance measured. To determine tolerance to Vincristine or Taxol cells were first subjected to 48h of doxycycline-induced Wapl depletion then exposed to repetitive cycles of Vincristine or Taxol treatment for 48h, followed by recovery in drug-free medium for 48h, where drug concentration was doubled every cycle. Concentrations ranged from 1nM to 3nM (A549) or 12nM (PC9 and H1299) of Taxol, and 4nM to 8nM (H1299) or 32nM (PC9 and A549) of Vincristine. Cells were maintained sub-confluent and split, as needed, during recovery days. Following final drug concentration cells were stained and analyzed as described above. Data presented represents average absorbances and standard deviation between three biological replicates. A students' 2-tailed t-test was used to determine statistical significance.

**EGFR sequencing**

Total RNA was isolated from primary mouse tumors using Trizol (Invitrogen). cDNA was transcribed with Superscript First-Strand Synthesis System (Invitrogen) and used as a template for subsequent PCR based studies. EGFR was amplified by PCR and products were cloned into a TOPO TA cloning vector (Invitrogen), transformed into bacteria and the inserts from individual clones sequenced. Primers are listed in Supplemental Table 1.

| RNAi | |
|---|---|
| **Gene Name** | **Target Sequence** |
| **siWapl #1** | GGAGUAUAGUGCUCGGAAU |
| **siWapl #2** | GAGAGAUGUUUACGAGUUU |
| **siWapl #3** | CAAACAGUGAAUCGAGUAA |
| **siWapl #4** | CCAAAGAUACACGGGAUUA |
| **shWapl #1** | GAATGATTCCAATCGTAAATA |
| **shWapl #2** | AGCTACTTGATAGCATATAAA |
| **Inducible-shWapl** | TTCGTCATGCATTCGGCATTT |

| qPCR | |
|---|---|
| **Gene Name** | **Sequence** |
| **GAPDH Forward** | CTA GCT GGC CCG ATT TCT CC |
| **GAPDH Reverse** | CGC CCA ATA CGA CCA AAT CAG A |
| **WAPL Forward** | TGG TTT GAA ATT GGG CCT GTT C |
| **WAPL Reverse** | AAC GGA CTA CCC TTA GCA CAA |
| **Slug Forward** | CGA ACT GGA CAC ACA TAC AGT G |
| **Slug Reverse** | CTG AGG ATC TCT GGT TGT GGT |
| **Vimentin Forward** | AGT CCA CTG AGT ACC GGA GAC |
| **Vimentin Reverse** | CAT TTC ACG CAT CTG GCG TTC |
| **Aurora B Forward** | CAG TGG GAC ACC CGA CAT C |
| **Aurora B Reverse** | GTA CAC GTT TCC AAA CTT GCC |
| **Beta Actin Forward** | ACA CCT TCT ACA ATG AGC |
| **Beta Actin Reverse** | ACG TCA CAC TTC ATG ARG |

| PCR amplification | |
|---|---|
| **Gene Name** | **Sequence** |
| **EGFR Forward** | ACA TCG TTC GGA AGC GCA CGC TG |
| **EGFR Reverse** | GAT TCC AAT GCC ATC CAC TTG ATA GG |

**Supplemental Table 1: primer and RNAi construct sequences used in this study**

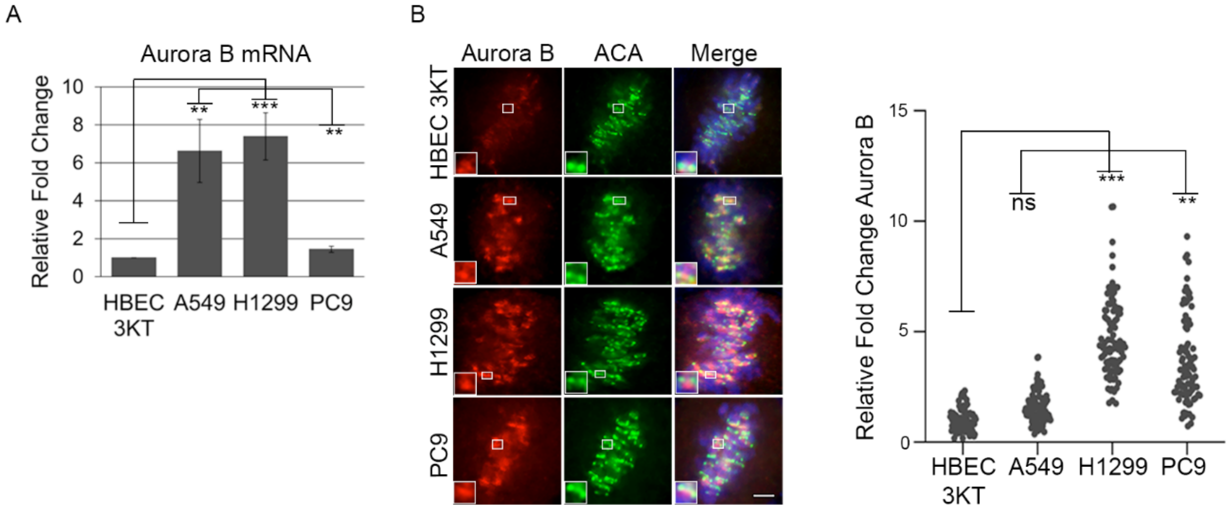


**Supplemental Figure 1: NSCLC cells display increased expression of Aurora B kinase** A) qPCR analysis of Aurora B mRNA levels in NSCLC cells (A549, H1299, PC9) compared to normal human bronchial epithelial cells (HBEC3-KT). B) Representation and quantification of centromere-localized Aurora B in NSCLC and HBEC3-KT cell lines. A minimum of 30 kinetochore pairs (3 per cell for 10 or more cells), for each of 3 biological replicates. Insets are of individual kinetochore pairs at 4x magnification. Scale bar is 5μm.

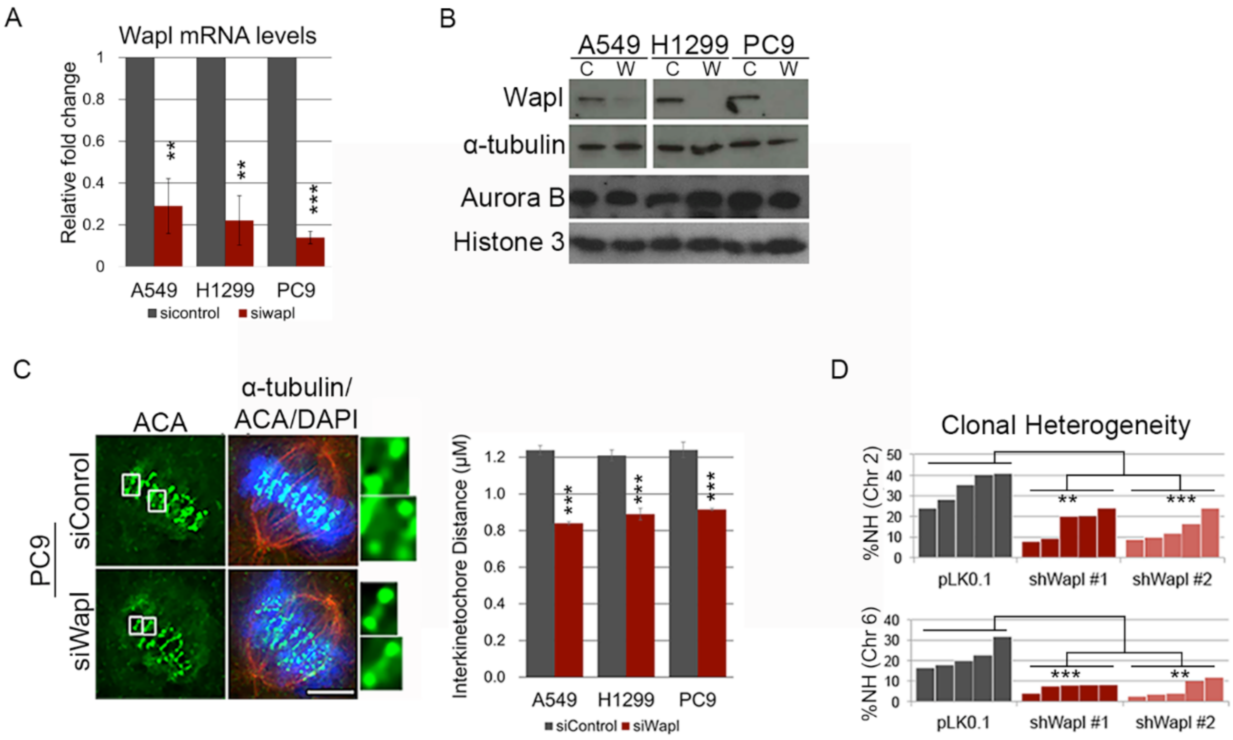


**Supplemental Figure 2: Increased cohesion reduces clonal heterogeneity in NSCLC cells** A) qPCR analysis of Wapl mRNA levels in control (Mock) and Wapl-depleted (Wapl KD) NSCLC cells. B) Western blot analysis of Aurora B and Wapl protein levels in control (C) and Wapl-depleted (W) NSCLC cell lines C) Representation of PC9 cells and quantification of inter-centromere distances in (A539, H1299, and PC9 cells) with and without Wapl depletion (siWapl and siControl, respectively). A minimum of 90 kinetochore pairs were measured (3/cell for 30 cells), for each of 3 biological replicates. Insets are of individual kinetochore pairs at 4x magnification. Scale bar is 5μm. D) Quantification of chromosome copy number heterogeneity (NH) in clonal populations of PC9 cells carrying an empty expression vector (pLKO.1) or one of two different shRNA constructs targeting Wapl for depletion (Wapl KD #1 and #2). Chromosome copy number was scored in a minimum of 300 cells for 5 clonal populations of each condition.

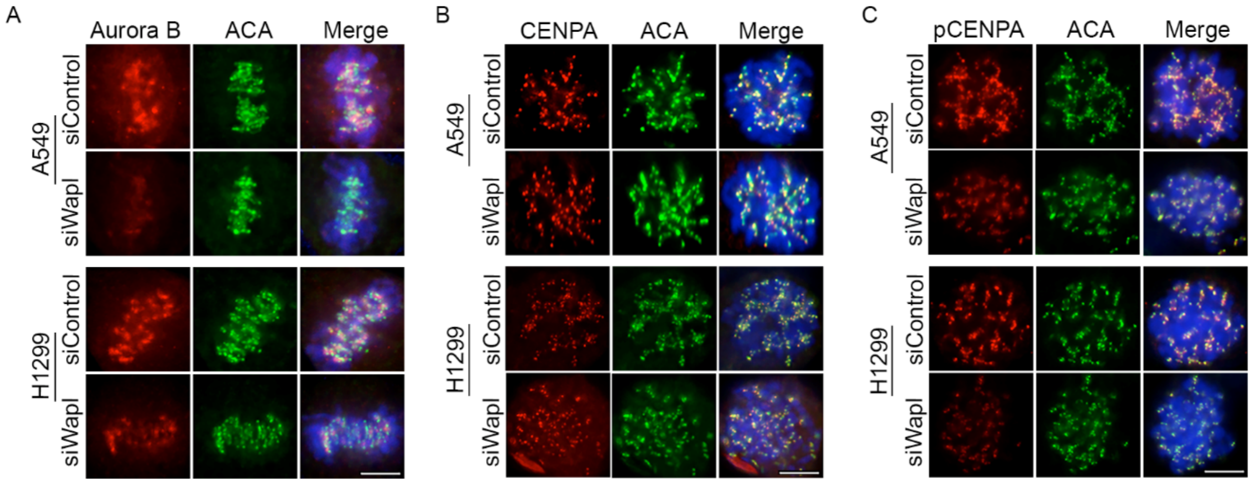


**Supplemental Figure 3: Enhanced cohesion limits Aurora B localization and activity at mitotic centromeres.** Representation and quantification of centromere-localized A) Aurora B, B) CENPA (total), and C) pCENPA (pSer7) following Wapl depletion (siWapl) or not (siControl) in A549 and H1299 cells. A minimum of 90 kinetochore pairs were measured (3/cell for 30 cells) for each of 3 biological replicates. Scale bar is 5μm.

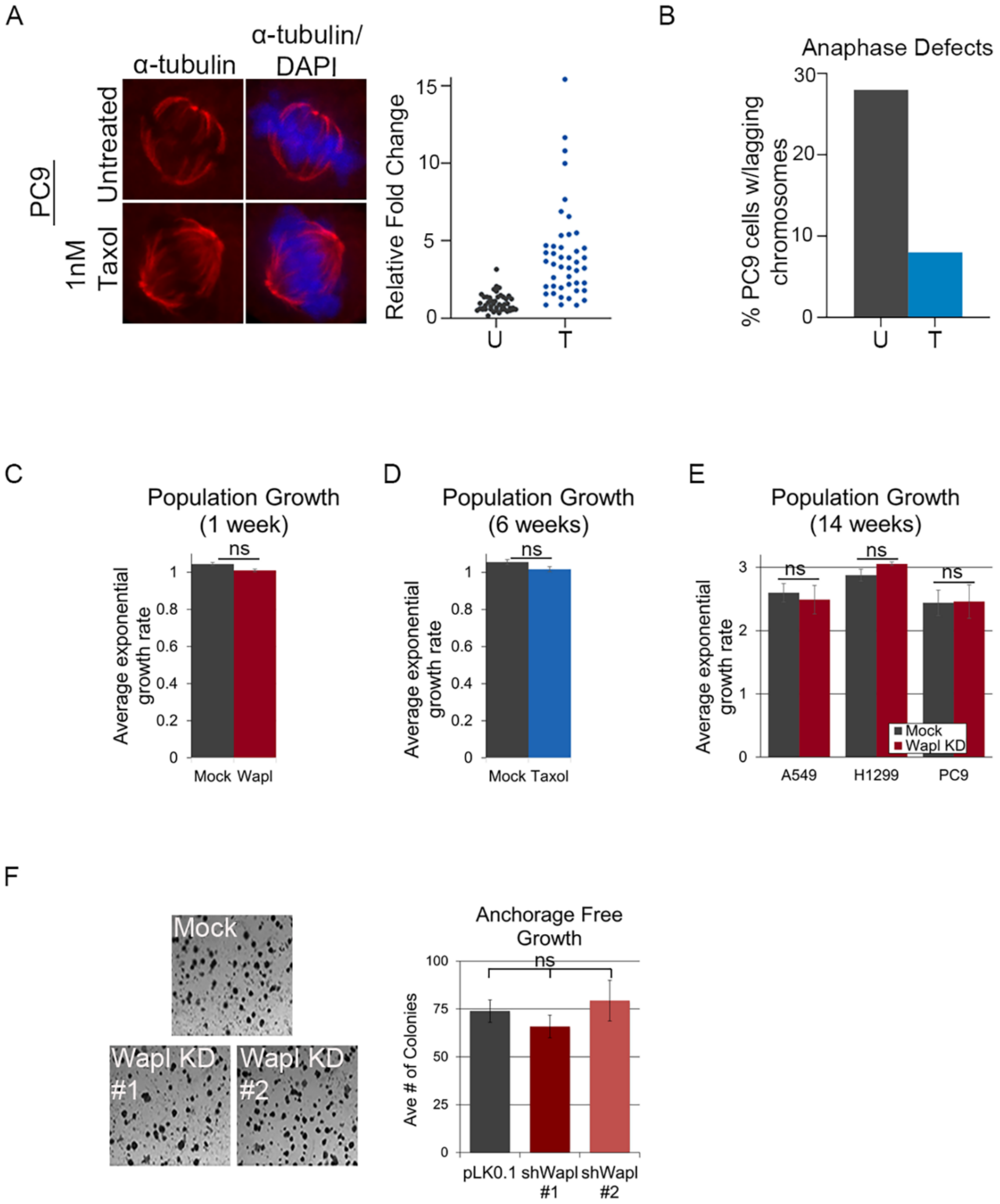


**Supplemental Figure 4: Suppression of nCIN limits does not compromise cell proliferation** A) Representation and quantification of cold stable microtubules in PC9 cells following acclimation to 1nM Taxol B) Quantification of anaphase lagging chromosomes in PC9 cells acclimated to 1nM Taxol. C-E) Average rate of population doublings following 1 week acute depletion of Wapl (C), 6 weeks of exposure to 1nM Taxol (D), or 14 weeks of constitutive Wapl depletion (E) in NSCLC cell lines as indicated. F) Representation and quantification of average colony number formed by control (PLK0.1) and Wapl depleted (Wapl KD#1 & #2) cell lines cultured under anchorage-free conditions.

A

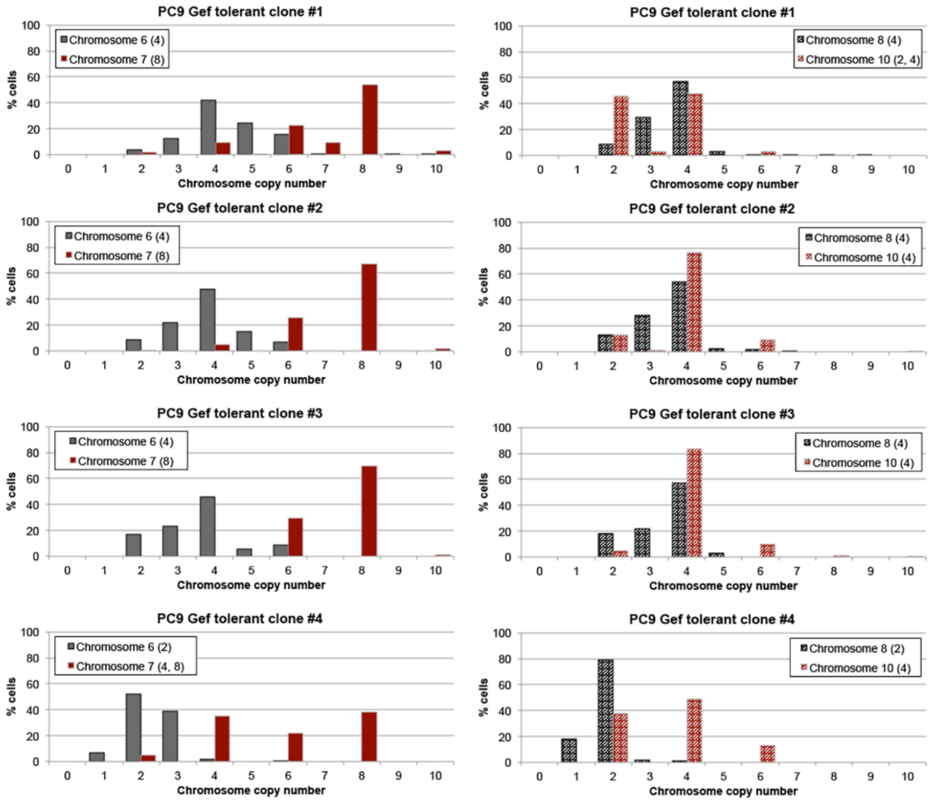
PC9 Gef tolerant clone #1
Chromosome 6 (4)
Chromosome 7 (8)
PC9 Gef tolerant clone #1
Chromosome 8 (4)
Chromosome 10 (2, 4)
PC9 Gef tolerant clone #2
Chromosome 6 (4)
Chromosome 7 (8)
PC9 Gef tolerant clone #2
Chromosome 8 (4)
Chromosome 10 (4)
PC9 Gef tolerant clone #3
Chromosome 6 (4)
Chromosome 7 (8)
PC9 Gef tolerant clone #3
Chromosome 8 (4)
Chromosome 10 (4)
PC9 Gef tolerant clone #4
Chromosome 6 (2)
Chromosome 7 (4, 8)
PC9 Gef tolerant clone #4
Chromosome 8 (2)
Chromosome 10 (4)
% cells
Chromosome copy number

B

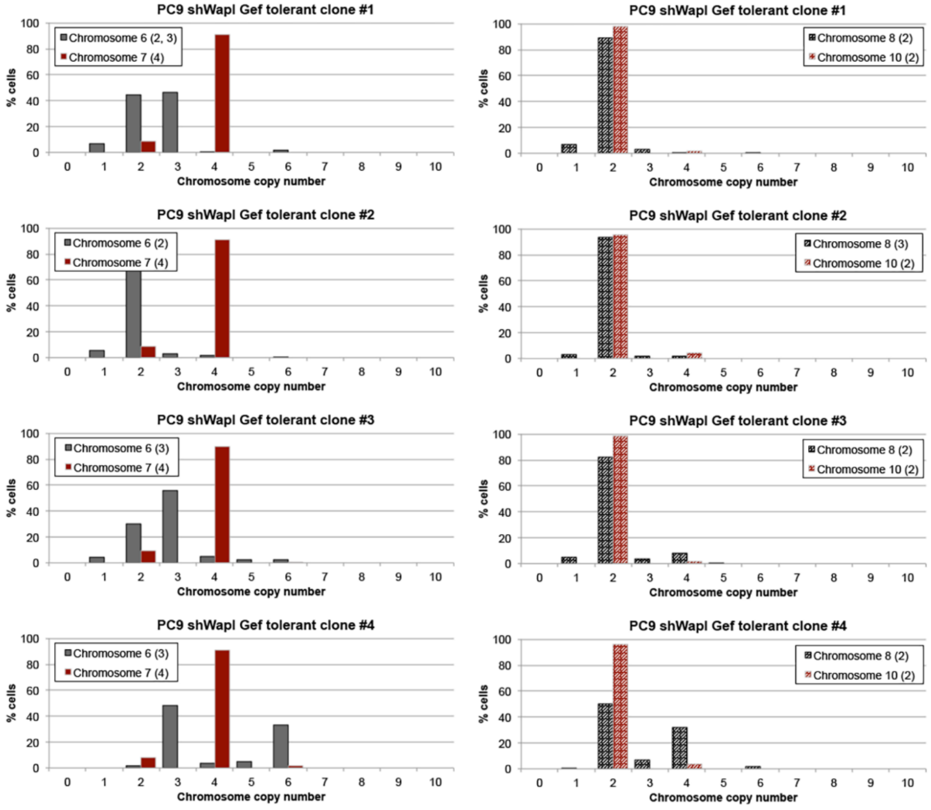
PC9 shWapl Gef tolerant clone #1
Chromosome 6 (2, 3)
Chromosome 7 (4)
PC9 shWapl Gef tolerant clone #1
Chromosome 8 (2)
Chromosome 10 (2)
PC9 shWapl Gef tolerant clone #2
Chromosome 6 (2)
Chromosome 7 (4)
PC9 shWapl Gef tolerant clone #2
Chromosome 8 (3)
Chromosome 10 (2)
PC9 shWapl Gef tolerant clone #3
Chromosome 6 (3)
Chromosome 7 (4)
PC9 shWapl Gef tolerant clone #3
Chromosome 8 (2)
Chromosome 10 (2)
PC9 shWapl Gef tolerant clone #4
Chromosome 6 (3)
Chromosome 7 (4)
PC9 shWapl Gef tolerant clone #4
Chromosome 8 (2)
Chromosome 10 (2)
% cells
Chromosome copy number

**Supplemental Figure 5: nCIN promotes acquisition of drug tolerance-promoting chromosome amplifications.** A & B) FISH analysis with centromere enumeration probes to measure chromosome copy number in individual cells for chromosomes 6, 7, 8 and 10 in Mock (A) or Wapl-depleted (B) Gefitinib-tolerant clones. Modal copy number for each chromosome is indicated in parentheses. Copy number of each chromosome probe was scored in a minimum of 300 cells for each cell line.

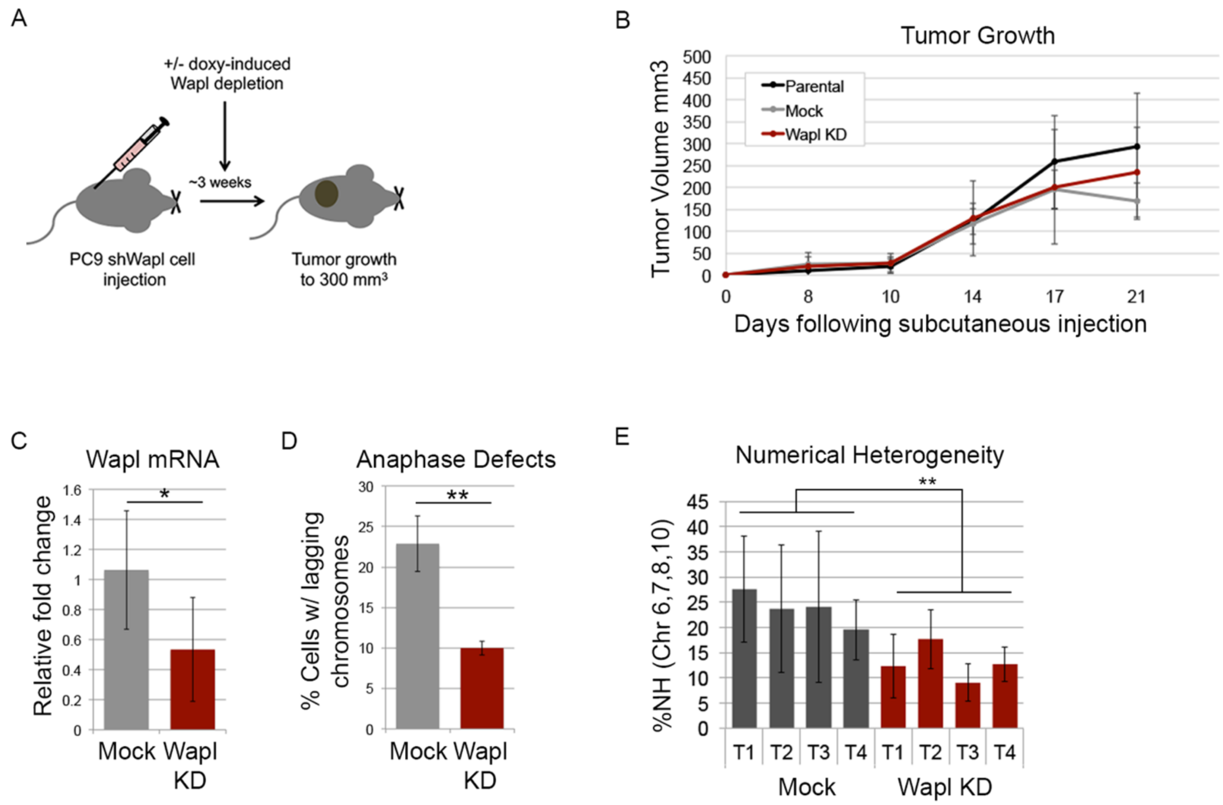


**Supplemental Figure 6: Wapl depletion is sufficient to suppress nCIN *in vivo***. A) Diagram of experimental set up: PC9 cells carrying an inducible shWapl construct were injected into the flanks of mice. Doxycycline was administered in the drinking water of one cohort to induce Wapl depletion and tumor initiation and growth were monitored. B) Measures of tumor volume, through duration of initial growth of tumors with (Wapl KD) or without (Parental, Mock) depletion of Wapl. C) qPCR analysis of Wapl mRNA levels in tumors at 3 weeks of growth. D) Quantification of anaphase lagging chromosomes in cells derived from mock and Wapl-depleted tumors. Graphs in B-D reflect average values and standard deviations from 4 independent tumor populations per condition. E) Tumor cells labelled with centromere enumeration FISH probes for chromosomes 6, 7, 8, and 10 and quantified for intratumor numerical heterogeneity.

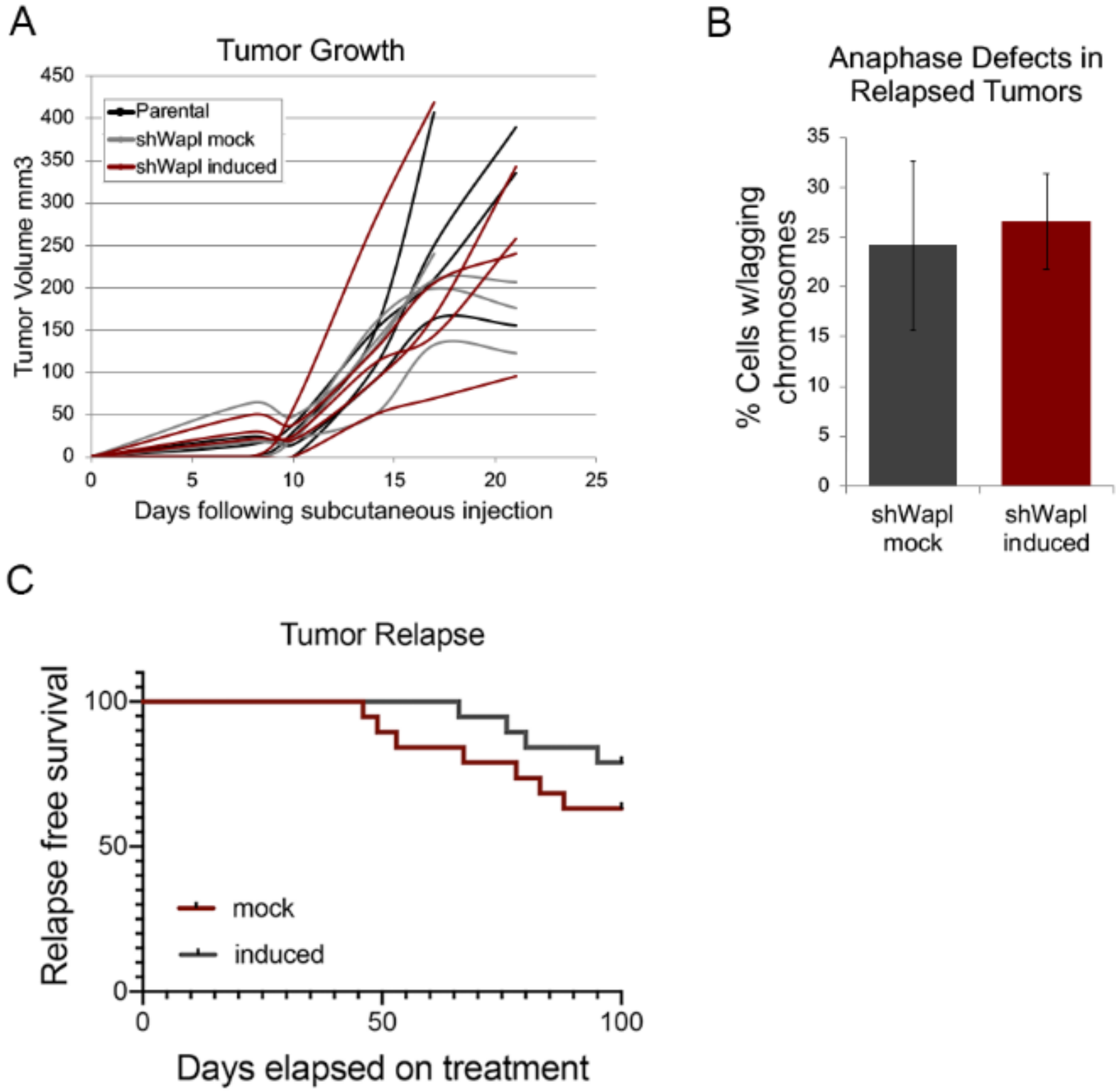


**Supplemental Figure 7: Suppression of CIN is not sustained in relapsed tumors.** A) Measures of individual tumor volumes over time, through duration of initial growth of drug naïve PC9 tumors with (Wapl KD) or without (Parental, Mock) depletion of Wapl. B) Quantification of anaphase defects in relapsed tumors arising from mock and Wapl-depleted PC9 cells. C) Relapse free survival of mice receiving a 5 + 2 regiment of Gefitinib with (Wapl KD) or without (Mock) concurrent induction of Wapl depletion.
